# Modulation of neural oscillations during working memory update, maintenance, and readout: an hdEEG study

**DOI:** 10.1101/2020.07.07.191619

**Authors:** Marianna Semprini, Gaia Bonassi, Federico Barban, Elisa Pelosin, Riccardo Iandolo, Michela Chiappalone, Dante Mantini, Laura Avanzino

## Abstract

Working memory (WM) performance is very often measured using the n-back task, in which the participant is presented with a sequence of stimuli, and required to indicate whether the current stimulus matches the one presented n steps earlier. In this study, we used high-density electroencephalography (hdEEG) coupled to source localization to obtain information on spatial distribution and temporal dynamics of neural oscillations associated with WM update, maintenance and readout. Specifically, we a priori selected regions from a large fronto-parietal network, including also the insula and the cerebellum, and we analyzed modulation of neural oscillations by event-related desynchronization and synchronization (ERD/ERS).

During update and readout, we found larger θ ERS and smaller β ERS respect to maintenance in all the selected areas. γ_LOW_ and γ_HIGH_ bands oscillations decreased in the frontal and insular cortices of the left hemisphere. In the maintenance phase we observed focally decreased θ oscillations and increased β oscillations (ERS) in most of the selected posterior areas and focally increased oscillations in γ_LOW_ and γ_HIGH_ bands in the frontal and insular cortices of the left hemisphere. Finally, during WM readout, we also found a focal modulation of the γ_LOW_ band in the left fusiform cortex and cerebellum, depending on the response trial type (true positive vs. true negative).

Overall, our study demonstrated specific spectral signatures associated with updating of memory information, WM maintenance and readout, with relatively high spatial resolution.

## Introduction

The n-back task—first described by Kirchner in 1958 (Kirchner, 1958)—is the most popular task used to measure working memory (WM), relying on the presentation of “rapidly, continuously changing information” to measure very short-term retention. In this task, participants are presented with a series of stimuli and are asked to indicate whether the current stimulus (probe) matches the stimulus presented n-stimuli back in the series. A recent review highlighted that WM at n-back is associated with a cerebral network that varies with stimulus type, presentation modalities and as a function of processing load (Mencarelli *et al*., 2019). Additionally, a number of evidence showed that specific frequency bands of electroencephalography (EEG) oscillations are of particular relevance for aspects of WM, such as the positive association between γ band activity (>40 Hz) and performance at higher WM loads in healthy populations (Crone *et al*., 2006; Lachaux *et al*., 2012; Roux *et al*., 2012; Honkanen *et al*., 2015; Kucewicz *et al*., 2017) and the association between θ oscillations and WM (Brookes *et al*., 2011; Burke *et al*., 2013; Hsieh & Ranganath, 2014). These observations have recently led to the use of non-invasive brain stimulation in combination with cognitive training for improving WM function (Hoy *et al*., 2015; Hill *et al*., 2019; Reinhart & Nguyen, 2019; Jones *et al*., 2020). Particularly, transcranial Alternating Current Stimulation (tACS) (Antal & Paulus, 2013; Helfrich *et al*., 2014) in the EEG range (conventionally: 0.1–80 Hz) in the frontal cortex is believed to directly modulate cortical oscillations and to impact sensory, perceptual and cognitive processes (Herrmann *et al*., 2013). However, to optimize such neuromodulation approach in cognitive rehabilitation of WM, we need a clear picture of the spatial distribution and temporal dynamics of cortical oscillations in the cerebral network involved in WM. In this context, high-density electroencephalography (hdEEG) provides us the possibility to gain information on the sources of the electrical oscillations underpinning cognitive processing with an optimal temporal resolution and an improved spatial resolution with respect to standard EEG (Michel et al., 2012). In particular, it is also fundamental to separately analyze oscillatory activity in the different phases of WM process: from the early phase of updating, i.e. the stored information at stimulus presentation, up to the usage of such information to guide action, going through the maintenance of information in face of other stimuli. Albeit it is not easy to disentangle the classic phases of working memory process (update, maintenance, readout) in the n-back task, in this study we aim to obtain information on spatial location and temporal dynamics of neural activity associated with the different phases. To this end, we used a custom developed pipeline for performing source localization from hdEEG data. This pipeline is able to detect multiple brain networks that are spatially similar to those obtained from fMRI data (Liu *et al*., 2017; Liu *et al*., 2018; Zhao *et al*., 2019).

We focused on correct update of stimuli by analyzing activity that was followed by a correct press n letters after (true positive) *and* activity that was followed by a correct no-press n letters after (true negative), with n being either 2 (2-back task) or 3 (3-back task). Furthermore, we analyzed hdEEG activity during the maintenance and readout of the n-back task. To analyze the maintenance phase, we observed the hdEEG activity in the single (2-back task) or in the two (3-back task) presented letters preceding the probe. Correct maintenance was identified when a correct response followed the appearance of the probe on the screen and when a correct no-response followed the appearance of the probe on the screen. Finally, hdEEG activity during the presentation of the probe was used to analyze the readout identified by probe letters correctly recognized as matching or non-matching the stimulus letter presented n-trials earlier.

## Materials and methods

### Data collection

We recruited 21 neurologically intact, right-handed subjects (9 females, age 30.9 ± 6.8 years, mean ± SD). All subjects provided written informed consent. The study conforms to the standard of the Declaration of Helsinki and was approved by the institutional ethical committee (CER Liguria Ref.1293 of September 12th, 2018).

The behavioral task consisted in a n-back working memory (WM) task (with n = 2, 3) as in (Hoy *et al*., 2015). Briefly, a series of random letters (A, B, C, D, E, F, G, H, I, O) was visually presented in sequence and the subject was required to respond with a button press when the currently presented letter corresponded to the letter presented n trials earlier (Figure 1 A). Each letter appeared on a screen for 500 ms with a 2000 ms delay between stimuli presentations (Figure 1 C).

**Figure 1.**
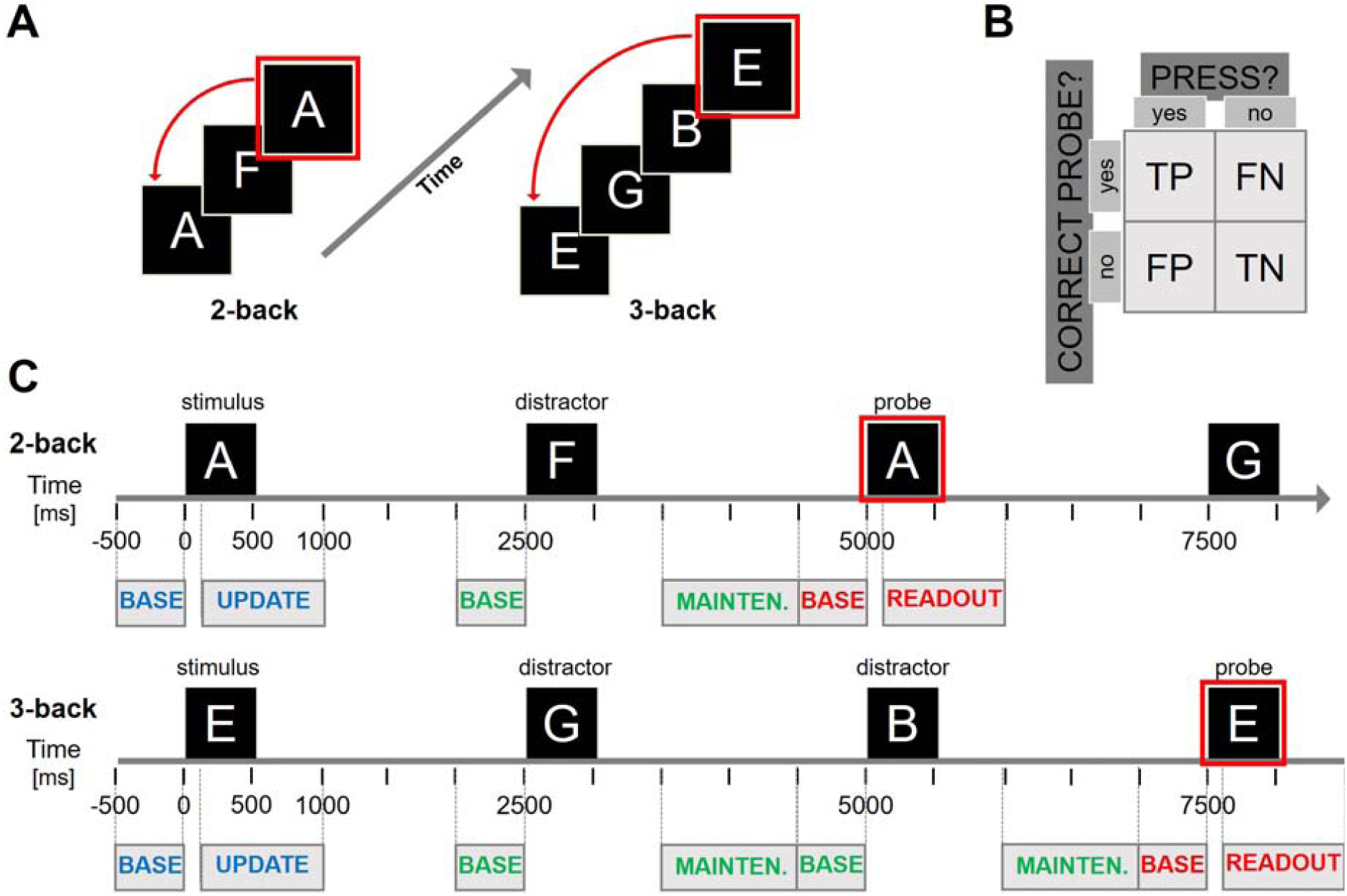
Outline of working memory task. **A)** Graphical representation of n-back tasks: the current letter (framed in red) must be compared with the one presented n times before, n being either 2 or 3 depending on the task. **B)** Contingency matrix of possible behavioral outcomes: true positive (TP) if a correct-probe letter was presented and the user correctly pressed the button; true negative (TN) if an incorrect-probe letter was presented and the users correctly did not press the button; false positive (FP) if an incorrect-probe letter was presented but the user pressed the button; false negative (FN) if a correct-probe letter was presented but the users did not press the button. **C)** Timeline representing 2-back (top) and 3-back (bottom) task timings and intervals chosen for hdEEG analysis. Letters appear on screen every 2500 ms and remain displayed for 500 ms. Analysis of memory *update* was performed by comparing baseline (500 ms preceding letter presentation, indicated as “BASE” in the figure) with a portion of signal ranging from 100 to 1000 ms post stimulus-letter onset (900 ms in total). Analysis of memory *update* was performed by comparing baseline (500 ms preceding distractor-letter presentation) with a portion of signal ranging from 1000 to 2000 ms post distractor-letter onset (1000 ms in total). Analysis of *readout* was performed by comparing baseline (500 ms preceding probe-letter presentation) with a portion of signal ranging from 100 to 600 ms post probe-probeletter onset (500 ms in total).

For hdEEG recording we used a 128 channel EEG recording system (actiCHamp, Brain Products), equipped with a trigger box handling external events. We collected hdEEG data at 1000 Hz sampling frequency, using the electrode FCz as physical reference. We also collected horizontal and vertical electrooculograms (EOG) from the right eye for further identification and removal of ocular-related artifacts.

The behavioral task was handled by a custom graphical user interface (GUI) developed in Matlab (The Mathworks). The GUI ran on a dedicated computer and was also responsible for sending task-related triggers to the EEG recording system. These triggers were sent through a NI USB board (National Instruments), which was also responsible of informing both the EEG recording system and the pc used for the cognitive task of a button press event.

### Cognitive processes underpinning working memory

With respect to the cognitive processes involved during the task, we distinguished between three WM phases: *update, maintenance* and *readout*. According to this distinction, the presented letters assume different roles. As an example, in Figure 1 A, for the 2-back case, the framed “A” represents the *probe* letter, the other “A” is the *stimulus* letter, and the “F” is the *distractor*; For the 3-back case, the framed “E” represents the *probe* letter, the other “E” is the *stimulus* letter, and “G” and “B” are *distractors*.

*Memory update* refers to the process of storing the presented letter (stimulus) for future comparison with the next probe letter (probe being either 2 or 3 trials later, depending on *n* value).

*Memory maintenance* refers to the process of keeping the previously presented letter in memory, when other letters (distractors) are presented before the probe letter (there is one distractor letter in the 2-back case, and two distractor letters in the 3-back case).

*Readout* corresponds to the processing of a behavioral response after the probe letter has been presented.

As depicted in Figure 1 B, by observing the behavioral responses we distinguished trials as belonging to one of the following categories:

- True positive (TP): probe letter correctly recognized as matching the stimulus letter (button press);
- True negative (TN): probe letter correctly recognized as non-matching the stimulus letter (no button press);
- False positive (FP): probe letter incorrectly recognized as matching the stimulus letter (button press);
- False negative (FN): probe letter incorrectly recognized as non-matching the stimulus letter (no button press).

In this work, we only observed brain responses during the well-performed trials, i.e. TP and TN, because the number of the badly performed trials (FP and FN) was too small.

We observed task performance by computing reaction time, defined as the delay between probe letter onset and button press for TP trials only, and accuracy, defined as the ratio of TP trials over the total number of response (TP + TN + FP + FN).

### hdEEG pre-processing and source localization

For analysis of hdEEG data, we made use of a tailored analysis pipeline that was recently developed to reconstruct source of neural oscillations (Liu *et al*., 2017).

We first attenuated the power noise in the EEG channels by using a notch filter centered at 50 Hz. Then, we detected channels with low signal to noise ratio and we labeled them as “bad channels”. We defined a channel as “bad” if it resulted as an outlier with respect to: i) the Pearson correlation of the signal in the frequency band 1-80 Hz against all the signals from all the other channels; and/or ii) the noise variance estimated in the frequency band 200-250 Hz, where the EEG contribution can be considered as negligible. The threshold to define an outlier was set to mean ± 3 standard deviation of the values. The bad channels were interpolated by using information coming from the neighboring channels, as implemented in the FieldTrip toolbox (http://www.fieldtriptoolbox.org/). EEG signals were then band-pass filtered (1-80 Hz) with a FIR zero-phase distortion filter and downsampled at 250 Hz.

Biological artefacts were rejected using Independent Component Analysis (ICA). Independent Components (ICs) were estimated with a fast fixed-point ICA (FastICA) algorithm (Hyvarinen & Oja, 2000), as described in (Mantini *et al*., 2008). ICs were marked as bad if correlation with the power of the EOG signals was higher than 0.2. The time courses of the ICs classified as bad were reconstructed at the channel level and subtracted from the data. EEG signals were then re-referenced with a customized version of the Reference Electrode Standardization Technique (REST) (Yao, 2001; Yao *et al*., 2005; Mantini *et al*., 2008; Liu *et al*., 2015).

As in (Liu *et al*., 2017), we generated a volume conductor head model with 128 electrodes positioned over a T1-weighted MR anatomical template, as in (Liu *et al*., 2017). Then, we segmented 12 tissue classes: skin, eyes, muscle, fat, spongy bone, compact bone, gray matter, cerebellar gray matter, white matter, cerebellar white matter, cerebrospinal fluid and brainstem and we assigned them with characteristics conductivity values, as in (Haueisen *et al*., 1997). To create a numerical approximation of the volume conduction model and to calculate the leadfield matrix, we used the Simbio finite element method (FEM) implemented in FieldTrip. The leadfield matrix estimated the relationship between the measured scalp potentials and the dipoles corresponding to brain sources, which were constrained by a regular 6 mm grid spanning the cortical, subcortical and cerebellar gray matter.

Sources reconstruction was performed with the exact low-resolution brain electromagnetic tomography eLORETA (Pascual-Marqui *et al*., 2011) algorithm, using both the artifacts-free hdEEG signals and the head model conductor.

### ERS-ERD analysis

We chose to analyze a specific set of regions of interest (ROIs) in the brain, whose activation was previously found related to the n-back task (Mencarelli *et al*., 2019). Table 1 summarizes the observed ROIs.

**Table 1.**
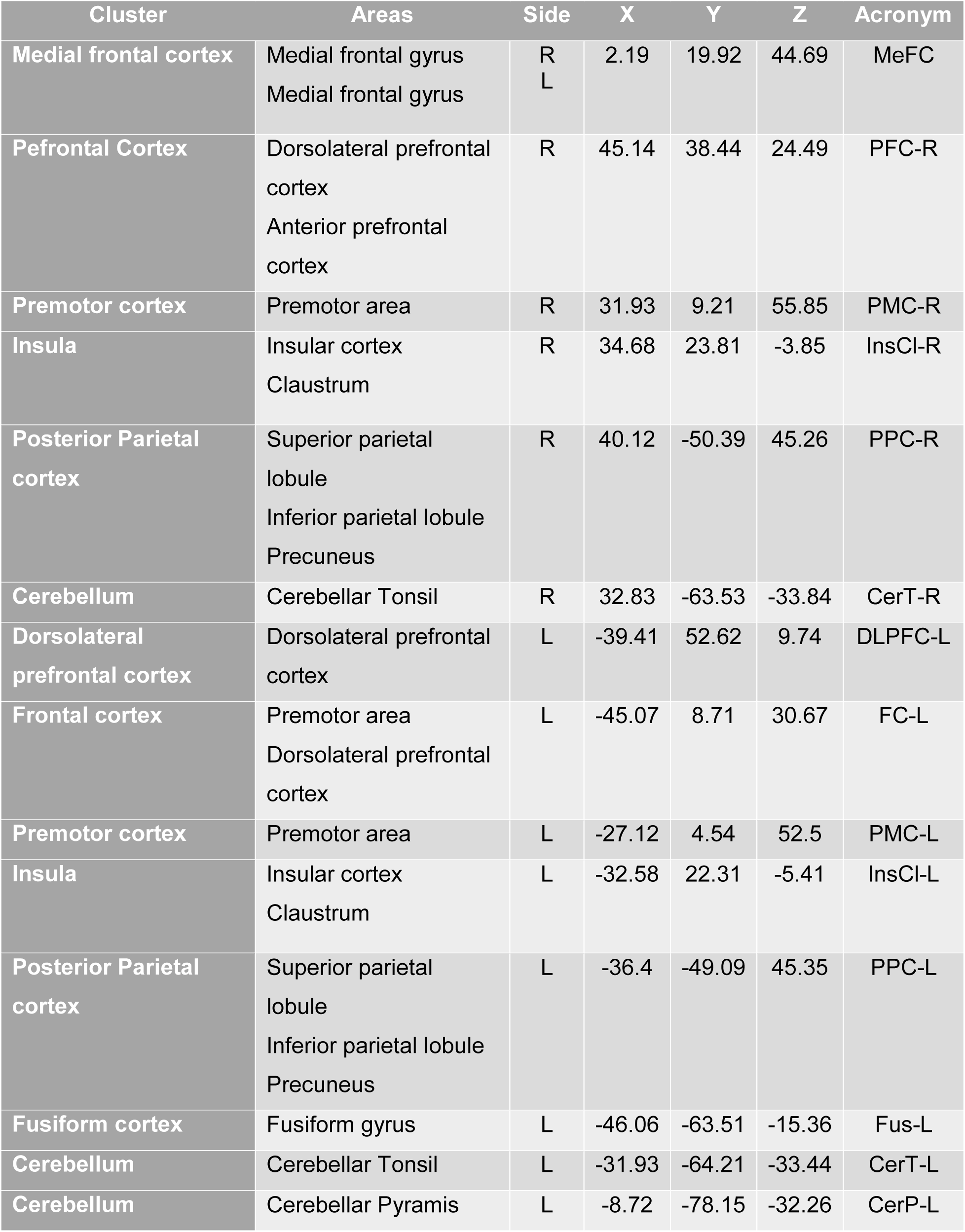
List of observed ROIs, areas they belong, and corresponding MNI coordinates.

We computed event related synchronization and desynchronization (ERS/ERD) of source reconstructed data filtered in different frequency bands and during different WM processing phases. Specifically, for each WM phase (update, maintenance and readout) we generated a spectrogram using Short-Time Fourier Transform for the frequency range 1–80 Hz, at steps of 1 Hz, and with temporal resolution equal to 100 ms. The spectrogram was epoched, according to each specific condition (see below) and then averaged. Finally, we calculated ERD/ERS intensity as the power change of the signal in a specific time range with respect to a reference period (baseline) (Pfurtscheller, 2001). We chose as baseline the 500 ms preceding letter presentation in all cases, as in (Hoy *et al*., 2016).

The observed frequency bands were θ (4-8 Hz), α (8-13 Hz), β (13-30 Hz), γ_LOW_ (30-50 Hz), and γ_HIGH_ (50-80 Hz). The δ band (1-4 Hz) was excluded from analysis, because it is often contaminated by motion artefacts.

Time range for *update* was set between 100 and 1000 ms post stimulus onset, similarly to (Hoy *et al*., 2016). The lower limit was set to 100 ms instead of 0 ms, because the visual system takes up to 150 ms to process visual stimuli (Thorpe *et al*., 1996).

Time range for *maintenance* was set between 1000 and 2000 ms post distractor onset. We chose this interval in order to discount the contribution provided by the update (100 – 1000 ms post letter presentation) of the distractors, which are, at the same time, probe letters for the following trials.

Time range for *readout* was set between 100 and 600 ms post stimulus onset. The lower limit was chosen as for the other trials in order to take into account the processing delays of the visual system (Thorpe *et al*., 1996), while for the upper limit the choice was data-driven and calculated according to the press distribution of all subjects during TP trials. Briefly, we grouped the press times of all subjects during all TP trials during 2-back (507 trials in total) and during 3-back (357 trials in total) and calculated the first percentile of each distribution (686 and 639 ms, respectively). We thus chose 600 ms as upper limit for both cases (2- and 3-back) and rejected trials where the press was made within 600 ms following letter presentation (in total we rejected 1 trial for 2-back and 7 trials for 3-back). The chosen temporal parameters are summarized in Table 2.

**Table 2.**
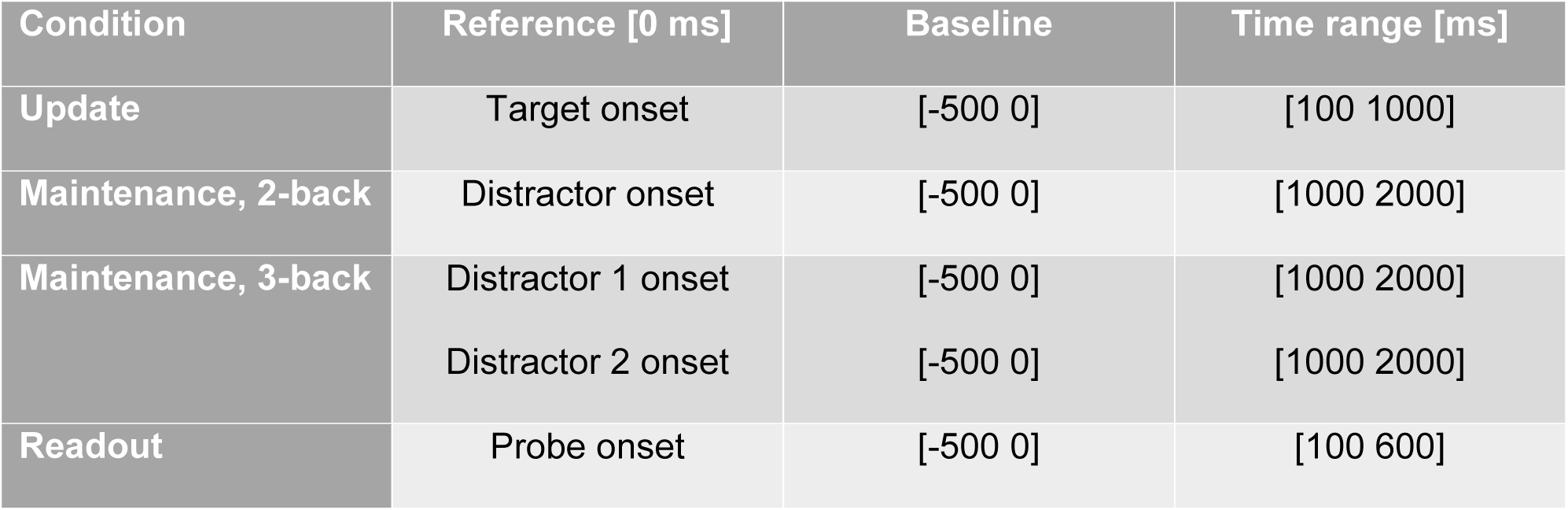
Temporal parameters used for ERS/ERD analysis of WM processing.

| Condition | Reference [0 ms] | Baseline | Time range [ms] |
| --- | --- | --- | --- |
| Update | Target onset | [-500 0] | [100 1000] |
| Maintenance, 2-back | Distractor onset | [-500 0] | [1000 2000] |
| Maintenance, 3-back | Distractor 1 onset | [-500 0] | [1000 2000] |
|  | Distractor 2 onset | [-500 0] | [1000 2000] |
| Readout | Probe onset | [-500 0] | [100 600] |

For statistical analysis of the data, we first assessed data normality with the one-sample Kolmogorov-Smirnov test. Then, a three-way repeated-measure analysis of variance (ANOVA) was run to test the influence on the mean ERD/ERS intensity on TASK (2-back and 3-back), PHASE (update, maintenance and readout) and TRIAL (TP, TN) as main factors within subjects, as well as of their interaction. This analysis was run separately for each ROI and for each frequency band. Post-hoc analysis was performed with Fisher Least Significant Difference method. The significance level was set to 0.05 for all analyses.

## Results

### Working Memory performance

Due to a lower WM load, best performance was obtained for the 2-back than the 3-back task. Figure 2 reports single subjects’ scores (panels A and D) as well as average scores (panels B and E) for the two tasks. We found a significant difference between accuracies obtained in the 2- and 3-back tasks (paired t-test, p = 0.0042) but not between reaction times (paired t-test, p = 0.21). In Figure 2 C we report the normalized distributions of reaction times for the 2-back (top) and 3-back (bottom) task. In Figure 2 F we plotted single subjects’ accuracy against mean reaction time for the two tasks. We found no correlation between these two measures.

**Figure 2.**
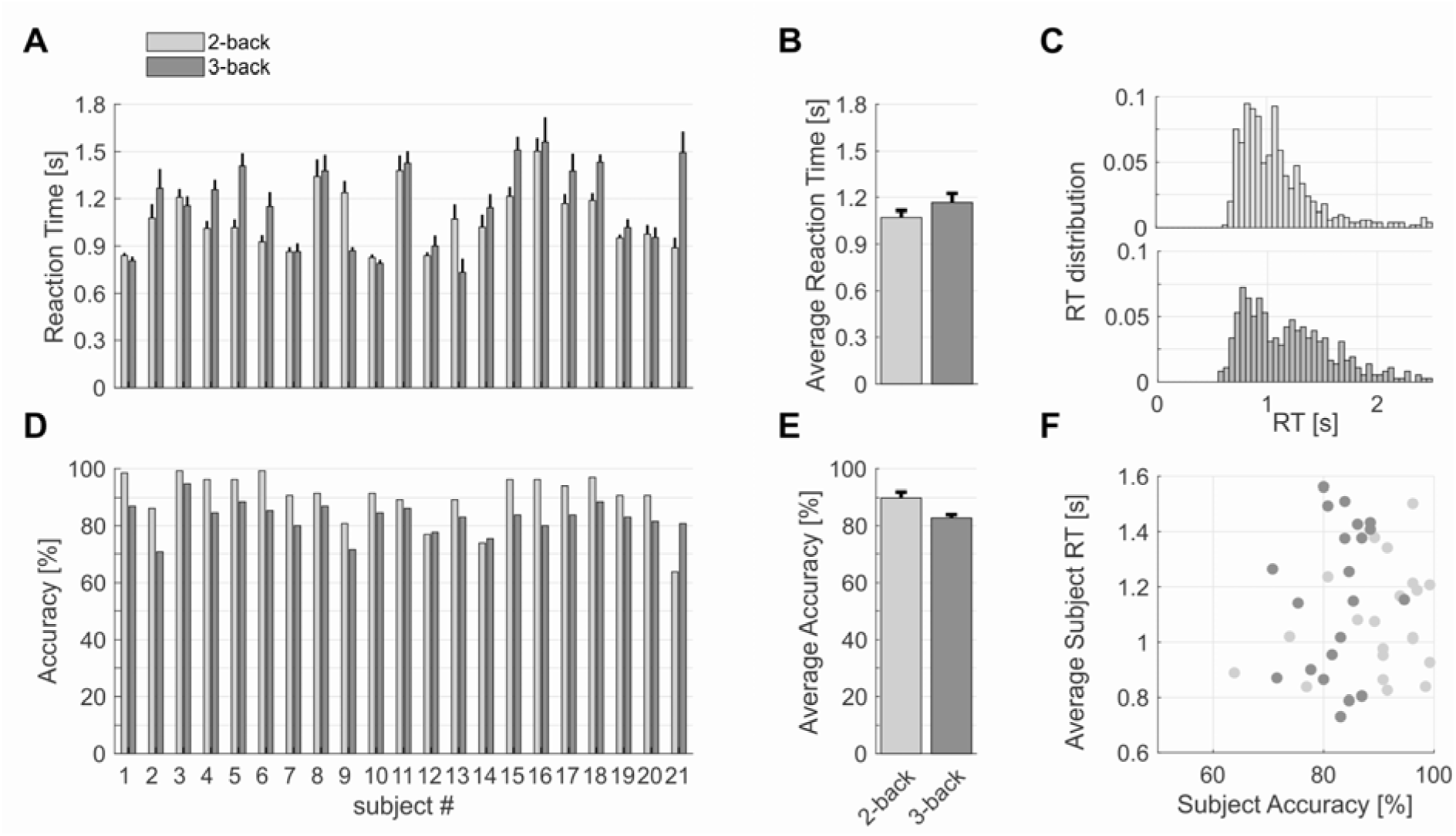
Cognitive performance of n-back task. **A)** Reaction times obtained by single subjects during the 2-back (light gray) and 3-back (dark gray) task (mean ± standard error of TP trials). **B)** Average reaction time obtained in the two tasks (mean ± standard error of single subjects’ scores). **C)** Normalized reaction times distribution of reaction times during the 2-back (top) and 3-back (bottom) tasks; data obtained from all the TP trials of all subjects. **D)** Accuracy obtained by single subjects during the 2-back (light gray) and 3-back (dark gray) task. **E)** Average reaction time obtained in the two tasks (mean ± standard error of single subjects’ scores). **F)** Accuracy vs mean reaction times of single subjects in the 2-back (light gray dots) and 3-back (dark gray dots) task.

### ERS/ERD analysis

For a specific set of ROIs (Table 1), we computed ERS/ERD for different tasks (i.e. 2- and 3-back), different trial types (i.e. TP and TN) and during the different phases of the working memory task (i.e. update, maintenance and readout).

We did not observe an effect of TASK on ERS/ERD modulation. Effect of TRIAL was observed in β, γ_LOW_ and γ_HIGH_ bands for few ROIs, with TP showing ERS and TN showing ERD (see Supplementary Materials). For most of the ROIs of interest, we observed a statistically significant effect of PHASE in all bands, with the exception of α. Significant interactions of main effects were never found between TASK and TRIAL, while in the other conditions (TASK*PHASE, TRIAL*PHASE and TASK*TRIAL*PHASE) were found for few ROIs in the γ_LOW_ band.

### Effect of PHASE

In the WM network, during update and readout, we found larger θ oscillations and smaller β oscillations respect to maintenance. In the maintenance phase we observed decreased θ oscillations with θ ERD in most of the selected posterior areas, and increased β oscillations (ERS).

Figure 3 reports ERS/ERD modulation in θ/β bands for all the ROIs of interest. A significant effect of PHASE was observed for the θ band in all the ROIs analyzed (Table 3 p ≤ 10^−5^ in all cases; Figure 3 panel A). Post-hoc analysis revealed an increase in θ oscillations in the update and readout with respect to maintenance, in all the analyzed ROIs (p ≤ 10^−3^ in all cases, except DLPFC-L, FC-L, and CerP-L with ≤ 10^−2^; Figure 3). For the β band, a significant effect of PHASE was also observed in all the ROIs (Table 3 p ≤ 0.001 in all cases, except PFC-R with p = 0.012, Figure 3 panel A). Post-hoc analysis revealed an increase in β oscillation in the maintenance and a decrease in β oscillation in the update and readout (maintenance vs update and readout, p≤ 10^−2^ in all cases).

**Table 3.** Results of ANOVA related to the main effect of PHASE. Asterisks report the level of significance (** p<0.01; * p<0.05).

| ROI | $\theta$ | $\beta$ | $\gamma_{\text{LOW}}$ | $\gamma_{\text{HIGH}}$ |
| --- | --- | --- | --- | --- |
| MeFC | $F_{2,40} = 21.40^{**}$<br>$p = 10^{-7}$ | $F_{2,40} = 14.61^{**}$<br>$p = 10^{-5}$ | $F_{2,40} = 0.07$<br>$p = 0.47$ | $F_{2,40} = 0.37$<br>$p = 0.69$ |
| DLPFC-R | $F_{2,40} = 16.08^{**}$<br>$p = 10^{-6}$ | $F_{2,40} = 4.91^{**}$<br>$p = 0.012$ | $F_{2,40} = 0.37$<br>$p = 0.68$ | $F_{2,40} = 0.47$<br>$p = 0.62$ |
| PMC-R | $F_{2,40} = 21.56^{**}$<br>$p = 10^{-7}$ | $F_{2,40} = 18.74^{**}$<br>$p = 10^{-6}$ | $F_{2,40} = 1.19$<br>$p = 0.31$ | $F_{2,40} = 0.33$<br>$p = 0.72$ |
| InsCI-R | $F_{2,40} = 18.57^{**}$<br>$p = 10^{-6}$ | $F_{2,40} = 7.67^{**}$<br>$p = 0.001$ | $F_{2,40} = 1.30$<br>$p = 0.28$ | $F_{2,40} = 1.81$<br>$p = 0.17$ |
| PPC-R | $F_{2,40} = 31.89^{**}$<br>$p = 10^{-9}$ | $F_{2,40} = 37.27^{**}$<br>$p = 10^{-10}$ | $F_{2,40} = 0.71$<br>$p = 0.49$ | $F_{2,40} = 1.57$<br>$p = 0.21$ |
| CerT-R | $F_{2,40} = 29.61^{**}$<br>$p = 10^{-8}$ | $F_{2,40} = 22.68^{**}$<br>$p = 10^{-7}$ | $F_{2,40} = 2.82$<br>$p = 0.07$ | $F_{2,40} = 4.42^*$<br>$p = 0.01$ |
| DLPFC-L | $F_{2,40} = 11.71^{**}$<br>$p = 10^{-5}$ | $F_{2,40} = 8.82^{**}$<br>$p = 0.0006$ | $F_{2,40} = 3.11^*$<br>$p = 0.048$ | $F_{2,40} = 3.16^*$<br>$p = 0.048$ |
| FC-L | $F_{2,40} = 13.55^{**}$<br>$p = 10^{-5}$ | $F_{2,40} = 10.11^{**}$<br>$p = 0.0002$ | $F_{2,40} = 4.43^{**}$<br>$p = 0.018$ | $F_{2,40} = 2.43$<br>$p = 0.10$ |
| PMC-L | $F_{2,40} = 2065^{**}$<br>$p = 10^{-7}$ | $F_{2,40} = 12.35^{**}$<br>$p = 10^{-5}$ | $F_{2,40} = 1.81$<br>$p = 0.17$ | $F_{2,40} = 0.95$<br>$p = 0.39$ |
| InsCI-L | $F_{2,40} = 16.68^{**}$<br>$p = 10^{-6}$ | $F_{2,40} = 10.63^{**}$<br>$p = 0.0001$ | $F_{2,40} = 5.44^{**}$<br>$p = 0.008$ | $F_{2,40} = 3.35^*$<br>$p = 0.045$ |
| PPC-L | $F_{2,40} = 28.54^{**}$<br>$p = 10^{-8}$ | $F_{2,40} = 23.11^{**}$<br>$p = 10^{-7}$ | $F_{2,40} = 0.98$<br>$p = 0.38$ | $F_{2,40} = 1.17$<br>$p = 0.32$ |
| Fus-L | $F_{2,40} = 27.72^{**}$<br>$p = 10^{-8}$ | $F_{2,40} = 13.34^{**}$<br>$p = 10^{-5}$ | $F_{2,40} = 5.78^{**}$<br>$p = 0.006$ | $F_{2,40} = 2.17$<br>$p = 0.12$ |
| CerT-L | $F_{2,40} = 24.78^{**}$<br>$p = 10^{-8}$ | $F_{2,40} = 15.58^{**}$<br>$p = 10^{-6}$ | $F_{2,40} = 4.68^*$<br>$p = 0.014$ | $F_{2,40} = 3.23^*$<br>$p = 0.049$ |
| CerP-L | $F_{2,40} = 13.33^{**}$<br>$p = 10^{-5}$ | $F_{2,40} = 15.03^{**}$<br>$p = 10^{-5}$ | $F_{2,40} = 2.24$<br>$p = 0.11$ | $F_{2,40} = 2.79$<br>$p = 0.07$ |

**Figure 3.**
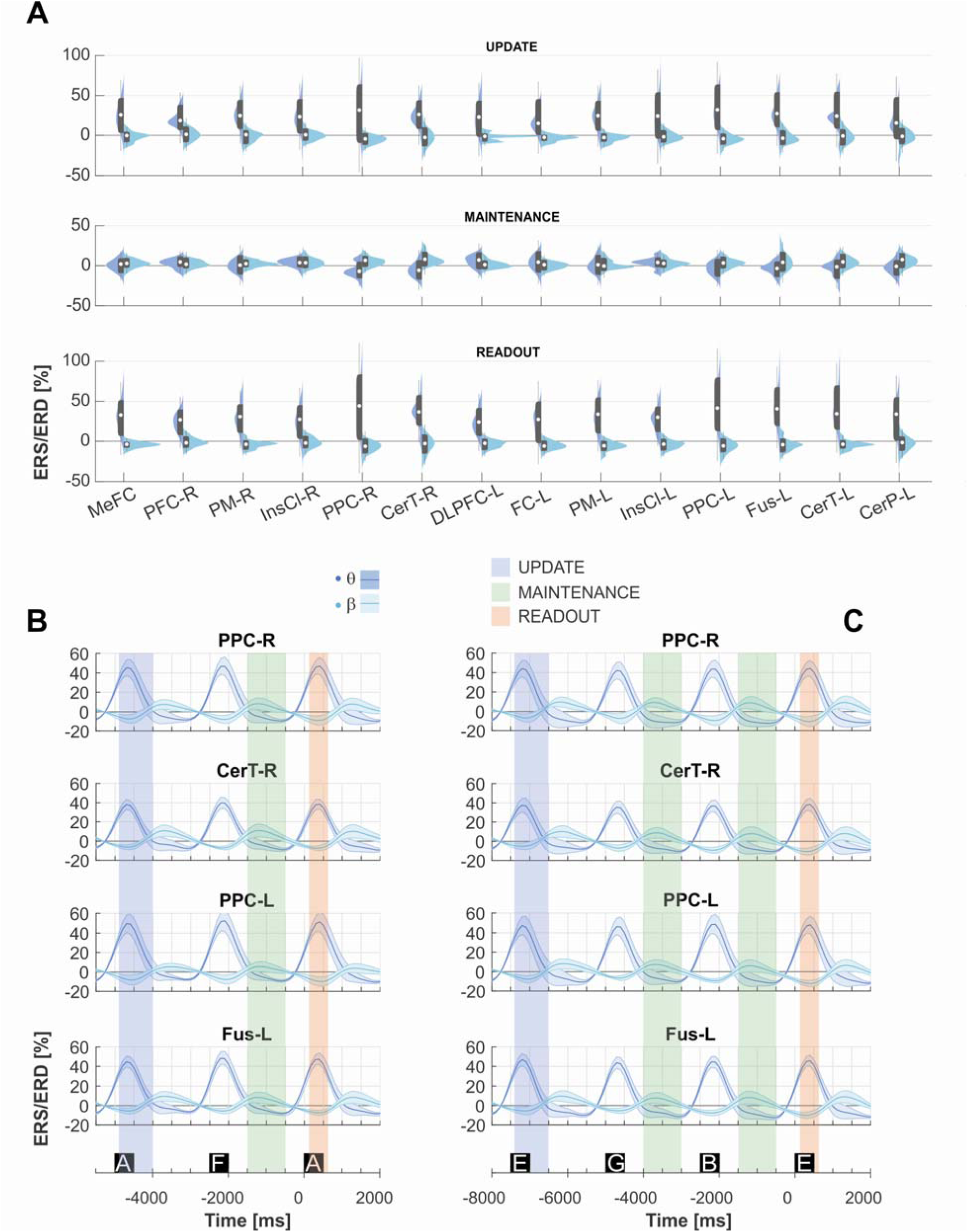
Effect of PHASE in the θ and β band. A) Violin plots of ERS/ERD variation in the θ (blue) and β (light blue) bands during update (top), maintenance (middle) and readout (bottom). Superimposed in grey are boxplots describing the median value (white dot), 25^th^ and 75^th^ percentiles (extremes of the thick grey line), and full data range (extremes of the thin grey line) of the distributions. B) Temporal evolution of band power in the θ (blue) and β (light blue) during the 2-back task for PPC-R, CerT-R, PPC-L, and Fus-L.). C) Temporal evolution of band power in the θ (blue) and β (light blue) during the 3-back task for PPC-R, CerT-R, PPC-L, and Fus-L.

For γ_LOW_ and γ_HIGH_ bands, ERS/ERD modulation was focally modified in the left hemisphere, in the insular, frontal cortex and in the cerebellar ROIs. Analogously to β activity, we observed in the insular and frontal cortex, smaller oscillations in the update and the readout with respect to maintenance. ERS/ERD modulation in γ_LOW_/γ_HIGH_ bands is reported in Figure 4 for the ROIs in which the effect of PHASE was found significant, and in Figure S1 for the other ROIs. Indeed, a significant effect of PHASE was observed mostly in the left hemisphere (Figure 4 panel A): γ_LOW_ oscillations were found significant (Table 3, p ≤ 0.05) in DLPFC-L, FC-L, IncCl-L, Fus-L, and CerT-L, while γ_HIGH_ oscillations in CerT-R, DLPFC-L, IncCl-L, and CerTL. Post-hoc analysis revealed a stronger increase in γ oscillation (ERS) in the maintenance phase than in update and readout (maintenance vs update and readout, p ≤ 0.04, Figure 4).

**Figure 2.**
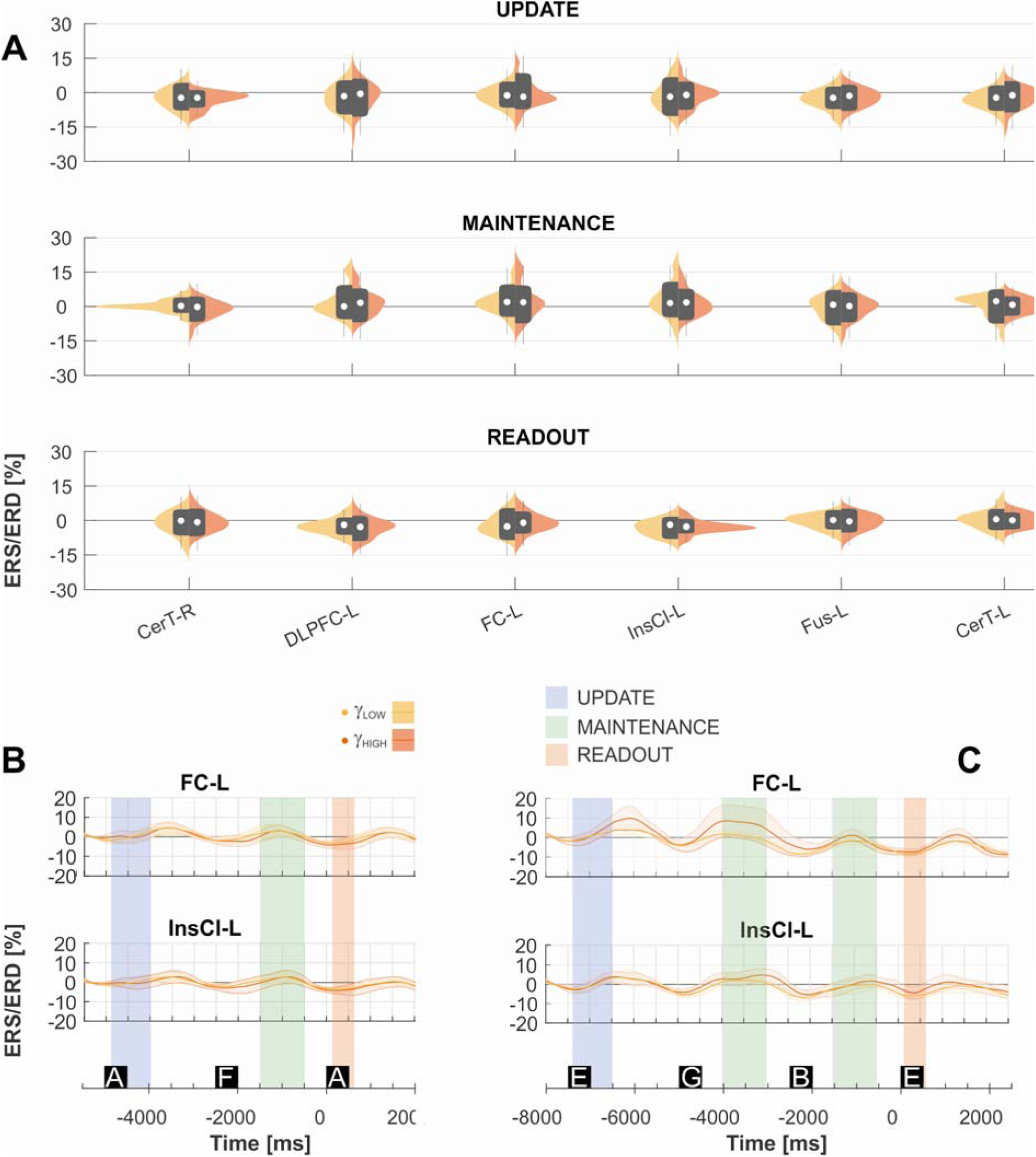
Effect of PHASE in the γ_LOW_ and γ_HIGH_ band. A) Violin plots of ERS/ERD variation in the γ_LOW_ (yellow) and γ_HIGH_ (orange) bands during update (top), maintenance (middle) and readout (bottom). Superimposed in grey are boxplots describing the median value (white dot), 25^th^ and 75^th^ percentiles (extremes of the thick grey line), and full data range (extremes of the thin grey line) of the distributions. B) Temporal evolution of band power in the γ_LOW_ (yellow) and γ_HIGH_ (orange) during the 2-back task for FC-L and InsCl-L.). C) Temporal evolution of band power in the γ_LOW_ (yellow) and γ_HIGH_ (orange) during the 3-back task for FC-L and InsCl-L.

### Interactions of main effects

Interactions of main effects were found only in the γ_LOW_ band. TASK*PHASE interaction was observed in cerebellum, in both hemispheres (Table 4; CerT-R p = 0.002, CerP-L p = 0.036), with update during 3-back showing a significant stronger ERD than during 2-back in the CerT-R (post-hoc analysis, p = 0.010) and a trend in the the CerP-L (post-hoc analysis, p = 0.09).

**Table 4.** Results of ANOVA related to the interaction of TASK*PHASE in the γ_LOW_ band. Asterisks report the level of significance (** p<0.01; * p<0.05; ^$^ trend). U, update; M, maintainance; R, readout. In the descriptive statistics, mean ± standard deviation is reported.

| ROI | Within test | Descriptive statistics | Post-hoc |
| --- | --- | --- | --- |
| CerT_R | $F_{2,40} = 7.13^{**}$ | U: 2-back ($-0.97 \pm 1.13$) 3-back ($-3.88 \pm 0.95$) | $p = 0.01^*$ |
| | $p = 0.002$ | M.: 2-back ($-0.84 \pm 1.16$) 3-back ($-0.86 \pm 1.12$) | $p = 0.25$ |
| | | R: 2-back ($-2.35 \pm 0.95$) 3-back ($-0.39 \pm 1.39$) | $p = 0.16$ |
| CerP_L | $F_{2,40} = 3.58^*$ | U: 2-back ($-0.32 \pm 1.20$) 3-back ($-2.84 \pm 0.82$) | $p = 0.09^{\$}$ |
| | $p = 0.036$ | M.: 2-back ($0.79 \pm 1.75$) 3-back ($0.05 \pm 1.20$) | $p = 0.09^{\$}$ |
| | | R: 2-back ($-0.80 \pm 0.91$) 3-back ($1.12 \pm 1.28$) | $p = 0.7$ |

In the left fusiform cortex and left cerebellum it was also observed a significant interaction TRIAL*PHASE (Table 5; CerT-L p = 0.013, Fus-L p = 0.002). Here we only expect interactions with the response phase, as trial type should not influence update and maintenance phases. Indeed, post-hoc analysis was found significant for CerT-L (p < 0.0005) and Fus-L (p = 0.0012), with γ oscillations increasing during readout of TP trials and decreasing during readout of TN trials. Finally, a significant TASK*TRIAL*PHASE interaction was also observed in the left fusiform cortex and right cerebellum (Table 6; CerT-R p = 0.017, Fus-L p = 0.03). Post-hoc analysis showed, for TP trials in both areas, stronger ERD for the 3-back task with respect to the 2-back task in the update phase (p = 0.006 for CerT-R and p = 0.014 for Fus-L). In CerT-R we also found a stronger ERS in the 3-back task with respect to the 2-back in the readout phase of TP trials (p = 0.048).

**Table 5.** Results of ANOVA related to the interaction of interaction of TRIAL*PHASE in the γ_LOW_ band. Asterisks report the level of significance (** p<0.01; * p<0.05; $ trend). U, update; M, maintainance; R, readout. TP, true positive trials; TN, true negative trials. In the descriptive statistics, mean ± standard deviation is reported.

| ROI | Within test | Descriptive statistics | Post-hoc |
| --- | --- | --- | --- |
| CerT_L | $F_{2,40} = 4.76^*$ | U: TP ($-3.42 \pm 1.26$) TN ($-1.88 \pm 0.88$) | $p = 0.2$ |
| | $p = 0.013$ | M: TP ($0.84 \pm 1.86$) TN ($-0.87 \pm 0.55$) | $p = 0.3$ |
| | | R: TP ($2.91 \pm 1.24$) TN ($-2.44 \pm 0.82$) | $p = 0.0005^{**}$ |
| Fus_L | $F_{2,40} = 6.96^{**}$ | U: TP ($-4.40 \pm 1.42$) TN ($-1.15 \pm 0.80$) | $p = 0.03^*$ |
| | $p = 0.002$ | M: TP ($0.41 \pm 1.83$) TN ($-0.53 \pm 0.64$) | $p = 0.9$ |
| | | R: TP ($3.13 \pm 1.42$) TN ($-1.98 \pm 0.75$) | $p = 0.0012^{**}$ |

**Table 6.** Results of ANOVA related to interaction of TRIAL*PHASE*TASK in the γ_LOW_ band. Asterisks report the level of significance (** p<0.01; * p<0.05; $ trend). U, update; M, maintainance; R, readout. TP, true positive trials; TN, true negative trials. In the descriptive statistics, mean ± standard deviation is reported.

| ROI | Within test | Descriptive statistics | Post-hoc |
| --- | --- | --- | --- |
| CerT-R | $F_{2,40} = 4.47^*$<br>$p = 0.017$ | TP:<br>U: 2-back($0.71 \pm 2.07$) 3-back ($-5.36 \pm 1.91$)<br>M: 2 back ($0.98 \pm 2.27$) 3-back ($-0.27 \pm 1.73$)<br>R.: 2-back($-1.74 \pm 1.54$) 3-back ($2.09 \pm 2.01$)<br>TN:<br>U: 2-back($-2.65 \pm 0.8$) 3-back ($-2.39 \pm 1.02$)<br>M.: 2 back ($0.69 \pm 0.94$) 3-back ($-1.4 \pm 1.03$)<br>R: 2-back ($-2.94 \pm 0.78$) 3-back ($-2.87 \pm 1.03$) | $p = 0.006^{**}$<br>$p = 0.64$<br>$p = 0.048^*$<br>$p = 0.80$<br>$p = 0.11$<br>$p = 0.95$ |
| Fus-L | $F_{2,40} = 3.70^*$<br>$p = 0.03$ | TP:<br>U: 2-back ($-0.29 \pm 1.81$) 3-back ($-8.5 \pm 2.32$)<br>M: 2-back ($-0.96 \pm 2.61$) 3-back ($0.14 \pm 2.05$)<br>R: 2-back ($3.28 \pm 1.62$) 3-back ($2.98 \pm 1.93$)<br>TN:<br>U: 2-back ($-0.84 \pm 1.08$) 3-back ($1.44 \pm 1.00$)<br>M: 2-back ($0.26 \pm 0.85$) 3-back ($1.32 \pm 0.78$)<br>R: 2-back ($-1.29 \pm 0.92$) 3-back ($-2.66 \pm 0.95$) | $p = 0.014^*$<br>$p = 0.71$<br>$p = 0.89$<br>$p = 0.66$<br>$p = 0.13$<br>$p = 0.24$ |

## Discussion

The main goal of this study was to obtain information on spatial location and temporal dynamics of neural oscillations associated with the different phases of working memory (update, maintenance, readout) in the n-back task.

Indeed, working memory is a high cognitive function that refers to the ability to encode, manipulate and retrieve information online and over a limited period of time (Baddeley, 1996). To this aim, taking advantage of recent developments on the accurate reconstruction of neural activity in the brain from hdEEG (Liu *et al*., 2017; Zhao *et al*., 2019), we analyzed spectral signatures associated with updating of memory information, its maintenance and its readout when used to inform and guide behavior. We a priori selected a large fronto-parietal network (Mencarelli *et al*., 2019), including also the insula, involved in memory storage, and the cerebellum as subcortical area that has demonstrated to be involved in cognitive functions (Strick *et al*., 2009).

The main results of the present study were the following: (i) in the update and readout, larger θ oscillations accompanied by smaller β oscillations in most of the selected areas and decreased γ_LOW_ and γ_HIGH_ bands in the frontal and insular cortices of the left hemisphere; (ii) in the maintenance, focally decreased θ oscillations and increased β oscillation in posterior areas and increased oscillation in γ_LOW_ and γ_HIGH_ bands in the frontal and insular cortices of the left hemisphere; (iii) in the readout, focal modulation of γ_LOW_ band in the left cortex (fusiform) and left cerebellum depending on the response (ERS in the TP trials and ERD in the TN trials). Noteworthy, only for γ_LOW_ band we observed that some of the modulations in the update and readout of TP trials were stronger for the 3-back than the 2-back task, suggesting that the cognitive load is playing a role in its modulation.

### EEG oscillations in relation to working memory: the update phase

Several recent studies highlighted the role played by θ oscillations in working memory. Particularly, activity in this band has been related to increases in the amount of information to be retained, with θ modulations localized to frontal (Gevins *et al*., 1997; Jensen & Tesche, 2002), hippocampal (Tesche & Karhu, 2000), and parietal (Sarnthein *et al*., 1998) regions. One of the core functions attributed to θ band oscillations in the hippocampal system is the temporal integration of cell assemblies (Buzsaki & Moser, 2013). Although first demonstrated for the tracking of spatial positions, the same mechanism may also support the representation and consolidation of sequentially organized memory traces (Lisman & Jensen, 2013).

Our findings revealed synchronization in the θ band in all the areas belonging to WM network following stimulus presentation, consistent with long-range coordination of neuronal activity within WM network. Indeed, neural oscillations are thought to play a central role in coordinating neural activity both in local networks (Gray *et al*., 1989; Womelsdorf *et al*., 2007) and over longer distances (von Stein & Sarnthein, 2000). Particularly, δ and θ oscillatory regimens are characterized by long-range interactions (von Stein & Sarnthein, 2000) requiring communication among several different areas. Simultaneously with an increase of θ oscillations, we also found a decrease of β oscillations at stimulus presentation in parietal and frontal areas of both hemispheres. In a recent study adopting magnetoencephalography (MEG) θ and β/γ activity were assessed during the n-back and the Sternberg tasks (Brookes *et al*., 2011). Similarly to our results, the authors found increased frontline θ power together with decreased power in the β/γ on task initiation. These oscillatory power decreases were most prominent in the 20–40 Hz frequency band, even if modulation could be observed up to 80 Hz, implying a broad-band response (Brookes *et al*., 2011).

Our findings are also consistent with the literature suggesting that θ and β/γ oscillations are linked. Whilst amplitude modulation θ and β bands oscillations were seen in almost all areas of WM network, modulation in γ band oscillations were detectable specifically in the clusters containing frontal cortex and insula/claustrum of the left hemisphere.

### EEG oscillations in relation to working memory: the maintenance phase

Our results were consistent also in the subsequent phase of WM process; i.e. the maintenance, with increased γ activity in the same areas in which γ activity was modulated in the update. In addition to γ_LOW_ synchronization in the frontal cortex and insula/claustrum of the left hemisphere, in the maintenance phase we observed increased β oscillation (ERS) for most of the selected areas and, accordingly to what observed in the update phase, decreased θ activity, specifically in the posterior areas.

In human MEG and EEG recordings, the maintenance of visual information in WM is associated with increased β and γ frequency band amplitudes (Tallon-Baudry *et al*., 1998; Osipova *et al*., 2006; Jokisch & Jensen, 2007; Haenschel *et al*., 2009; Palva *et al*., 2011). Related to β oscillations, although β has been widely studied for movement, it has also been suggested a role in cognitive functions such as WM (Lundqvist *et al*., 2011; Lundqvist *et al*., 2016; Lundqvist *et al*., 2018). Recent studies recorded prefrontal activity in monkeys performing a delayed match-to-sample task, in which several objects had to be encoded, maintained, and tested sequentially over several seconds (Lundqvist *et al*., 2016). During encoding, brief γ bursts were associated with spiking activity while β bursts were reduced. Then, in the following delay period, moderate increase of β was observed except at the very end, when information was needed again. At that point, β was reduced and γ increased. The authors speculated that the intermediate elevation of β during the delay period relative to the low levels seen at encoding and readout might serve to protect the current working memory contents from interference. Indeed, human studies have shown increases of prefrontal β when subjects must filter out distractors (Zavala *et al*., 2015; Zavala *et al*., 2017) or prevent encoding (Hanslmayr *et al*., 2014).

γ band oscillations have been suggested to represent a generic mechanism for the representation of individual WM items, irrespective of WM content and format (Roux & Uhlhaas, 2014). This is because the synchronization of neuronal discharges at γ frequencies supports the integration of neurons into cell assemblies in different cortical and subcortical structures (Singer, 2009) and thus could represent an effective representational format for WM information (Roux & Uhlhaas, 2014). γ modulations were observed focally in the prefrontal cortex and insula/claustrum of the left hemisphere. Insula is particularly involved in n-back tasks based on the visual presentation of numbers (Mencarelli *et al*., 2019), possibly linked to its phonological function (Chee *et al*., 2004). Visual WM operations may rely on activation of letter representations in insular cortex, via top-down feedback from neocortical areas including the prefrontal cortex (Moore *et al*., 2006). Thus, top-down input from the prefrontal cortex can additionally promote maintenance of visual images in the face of distraction (Sakai *et al*., 2002; Miller *et al*., 2018). Following this model, ERS in γ band in these areas may suggest critical role of this network in visual WM maintenance.

### EEG oscillations in relation to working memory: the readout phase

The temporal window for the readout started together with the update, 100 ms after letter presentation. Thus the overlap of θ/β oscillations in the readout (i.e.: increased θ oscillations and reduced β oscillations) with respect to update may be suggestive of the overlap of cognitive processes. Indeed in n-back task, it is difficult to disentangle between update and readout, since every new stimulus has to be encoded and simultaneously compared to the 2 or 3 stimuli preceding it, in order to be recognized and to trigger the correct response.

However, in addition to θ/β modulation, in the readout we also observed modulation in the γ_LOW_ band activity depending on readout process. Indeed, modulation of γ_LOW_ activity differed when decision was to press the button (TP) or not (TN). γ oscillations increased in the left fusiform cortex and left cerebellum when subjects had to decide that the probe letter was equal to the stimulus (TP trials) whereas γ oscillations decreased in the same areas when subjects had to decide that the probe letter differed from the stimulus (TN trials). In TP trials, increased γ oscillations in the readout phase are consistent with recent evidence coming from animal studies with local field potential recordings (Lundqvist *et al*., 2011; Lundqvist *et al*., 2016; Lundqvist *et al*., 2018), suggesting a role for γ oscillations when working memory needs to be read out. We also generalize this phenomenon to a process of readout instrumental to inform motor behavior (like a button press). The left fusiform gyrus has been connected with visual word processing (Cohen *et al*., 2002; Dehaene & Cohen, 2011; Price & Devlin, 2011; Wandell, 2011) and represents both phonological information in addition to orthographic information (Zhao *et al*., 2017). Cerebellar engagement in working memory tasks is reliably reported across multiple studies (Schmahmann & Pandya, 1997; Strick *et al*., 2009). Particularly, connections with association cortices (including the prefrontal cortex) are mainly located within posterior cerebellar lobules (including cerebellum tonsil and pyramis), which provide the anatomic substrate for cerebellar involvement in cognition. Taken together, we can suppose a network based on letter recognition, attention based motion processing and selection of WM information for action preparation, specifically active when the response to be selected is a motor output (and not to suppress the motor output, as it happens in the TN trials).

## Conclusions

Overall, our study demonstrated specific spectral signatures based on hdEEG associated with updating of memory information, working memory maintenance and readout, with relatively high spatial resolution. Considering that n-back task is largely used in clinical settings for both diagnosis and rehabilitation, our findings may support the targeted use of non-invasive neuromodulation techniques to boost the WM process in diseases.

## Acknowledgements

The authors gracefully acknowledge Martina Putzolu and Marta Carè for assistance during data acquisition.

