## supplementary material for "Modulation of neural oscillations during working memory update, maintenance, and readout: an hdEEG study"

### Effect of TRIAL

Table 1S shows the results of ANOVA for TRIAL as main effect. We found that TP and TN trials were respectively characterized by an increase and decrease of oscillations. This effect was found significant (p ≤ 0.04) in the β band for right hemisphere and both cerebellar ROIs, while in γ bands for left subcortical and cerebellar ROIs. We also found a significant effect (p = 0.035) for PPC-R in the γ_LOW_ band only.

**Figure 1S**

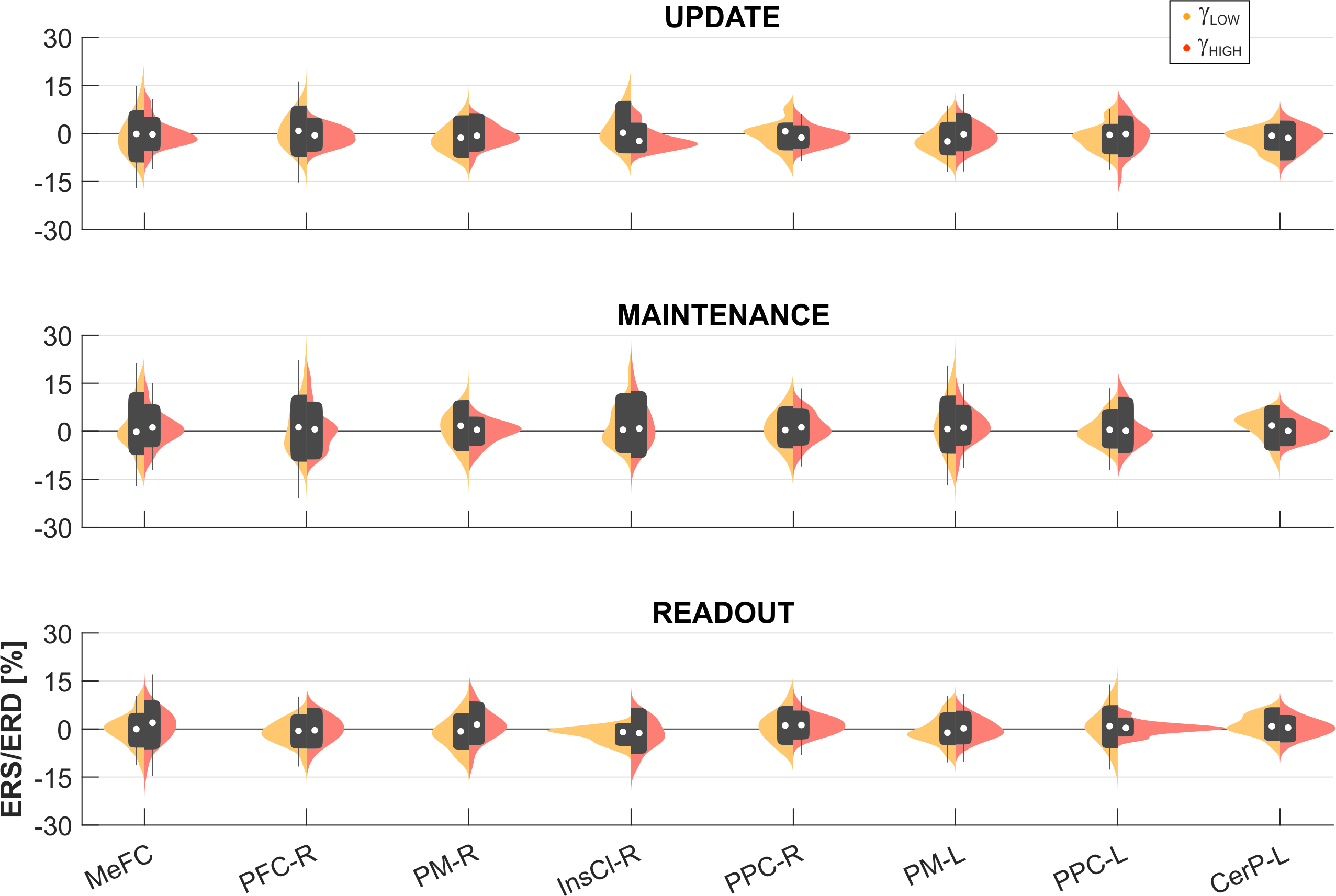

**Figure 1S:** Effect of PHASE in the γ_LOW_ and γ_HIGH_ band. Violin plots of ERS/ERD variation in the γ_LOW_ (yellow) and γ_HIGH_ (orange) bands during update (top), maintenance (middle) and readout (bottom). Superimposed in grey are boxplots describing the median value (white dot), 25th and 75th percentiles (extremes of the thick grey line), and full data range (extremes of the thin grey line) of the distributions.

**Table 1S:** results of ANOVA main effect of TRIAL. Asterisks report the level of significance (** p<0.01; * p<0.05).

| ROI | θ | β | γ_LOW_ | γ_HIGH_ |
| --- | --- | --- | --- | --- |
| MeFC | F_1.20_ = 0.038  p = 0.84 | F_1.20_ = 4.36*  p = 0.04 | F_1.20_ = 1.06  p = 0.31 | F_1.20_ = 0.009  p = 0.92 |
| PFC-R | F_1.20_ = 1.92  p = 0.18 | F_1.20_ = 14.24**  p = 0.001 | F_1.20_ = 1.47  p = 0.23 | F_1.20_ = 1.23  p = 0.28 |
| PMC-R | F_1.20_ = 1.92  p = 0.18 | F_1.20_ = 14.24**  p = 0.001 | F_1.20_ = 1.47  p = 0.23 | F_1.20_ = 1.23  p = 0.28 |
| InsCl-R | F_1.20_ = 1.00  p = 0.32 | F_1.20_ = 13.18**  p = 0.001 | F_1.20_ = 2.15  p = 0.15 | F_1.20_ = 0.35  p = 0.55 |
| PPC-R | F_1.20_ = 0.09  p = 0.76 | F_1.20_ = 2.47  p = 0.13 | F_1.20_ = 5.08  p = 0.035 | F_1.20_ = 3.26  p = 0.08 |
| CerT-R | F_1.20_ = 0.008  p = 0.92 | F_1.20_ = 7.63*  p = 0.01 | F_1.20_ = 1.81  p = 0.19 | F_1.20_ = 2.14  p = 0.15 |
| DLPFC-L | F_1.20_ = 0.27  p = 0.61 | F_1.20_ = 0.05  p = 0.81 | F_1.20_ = 0.009  p = 0.92 | F_1.20_ = 1.51  p = 0.23 |
| FC-L | F_1.20_ = 0.06  p = 0.79 | F_1.20_ = 0.06  p = 0.80 | F_1.20_ = 0.003  p = 0.95 | F_1.20_ = 0.04  p = 0.83 |
| PMC-L | F_1.20_ = 1.79  p = 0.19 | F_1.20_ = 0.28  p = 0.60 | F_1.20_ = 0.17  p = 0.68 | F_1.20_ = 0.0009  p = 0.97 |
| InsCl-L | F_1.20_ = 0.15  p = 0.70 | F_1.20_ = 3.84  p = 0.06 | F_1.20_ = 0.03  p = 0.86 | F_1.20_ = 2.47  p = 0.13 |
| PPC-L | F_1.20_ = 0.009  p = 0.93 | F_1.20_ = 2.60  p = 0.12 | F_1.20_ = 1.48  p = 0.23 | F_1.20_ = 3.25  p = 0.08 |
| Fus-L | F_1.20_ = 1.67  p = 0.21 | F_1.20_ = 1.35  p = 0.25 | F_1.20_ = 0.96  p = 0.33 | F_1.20_ = 5.91*  p = 0.024 |
| CerT-L | F_1.20_ = 0.35  p = 0.55 | F_1.20_ = 3.01  p = 0.097 | F_1.20_ = 7.56*  p = 0.012 | F_1.20_ = 6.93*  p = 0.015 |
| CerP-L | F_1.20_ = 0.033  p = 0.85 | F_1.20_ = 5.21*  p = 0.033 | F_1.20_ = 4.94*  p = 0.037 | F_1.20_ = 3.15  p = 0.09 |
